# Membrane interactions accelerate the self-aggregation of huntingtin exon 1 fragments in a polyglutamine length-dependent manner

**DOI:** 10.1101/2020.06.24.169060

**Authors:** A. Marquette, B. Bechinger

**Affiliations:** University of Strasbourg/CNRS; CNRS

**Keywords:** circular dichroism, DLS, thioflavin T fluorescence, peptide-lipid interactions, huntingtin, Huntington’s disease, amyloid, htt17, membrane-driven aggregation

## Abstract

The accumulation of aggregated protein is a typical hallmark of many human neurodegenerative disorders including Huntington’s disease. Misfolding of the amyloidogenic proteins gives rise to self-assembled complexes and fibers. The huntingtin protein is characterized by a segment of consecutive glutamines, which when exceeding a certain number of residues results in the occurrence of the disease. Furthermore, it has also been demonstrated that the 17-residue amino-terminal domain of the protein (htt17), located upstream of this polyglutamine tract, strongly correlates with aggregate formation and pathology. Here we demonstrate that membrane interactions strongly accelerate the oligomerization and β-amyloid fibril formation of htt17-polyglutamine segments. By using a combination of biophysical approaches the kinetics of fibre formation has been quantitatively investigated and found to be strongly dependent to the presence of lipids, the length of the polyQ expansion and the polypeptide-to-lipid ratio. Finally, the implications for therapeutic approaches are discussed.

**Statement of significance:** The quantitative analysis of the aggregation kinetics of amino-terminal fragments of huntingtin demonstrate the importance of the 17-residue amino-terminal membrane anchor and a resulting dominant effect of membranes in promoting the aggregation of polyglutamines. Other parameters further modulating the association kinetics are the length of the polyglutamine stretch and the peptide concentration. The findings can have important impact on finding new therapies to treat Huntington’s and other polyglutamine related diseases.

## INTRODUCTION

At least nine different hereditary diseases that are related to the expansion of a polyglutamine (polyQ) domain are known to date (1). These so-called CAG repeat pathologies are all related by the propensity of their associated polypeptide to form insoluble β-sheet rich amyloid fibrils. Polyglutamine expansion promotes the self-assembly of fibrils and other types of aggregates that accumulate in inclusions found for example in the brain tissues of patients. Using modern imaging techniques these have been described as dynamic phase separated gel-like structures or as coexisting liquid / solid condensates (2, 3). In the case of Huntington’s disease (HD) the age of development and the severity of the disease correlate with the length of the polyQ stretch which is located within the amino-terminal domain of huntingtin, a protein encompassing about 3500 amino acids (1, 4). Symptoms of the disease develop when the polyQ expansion exceeds a critical length of ∼ 37 glutamines. Although the genetics of HD are well studied, the exact biological functions of huntingtin remain speculative and the exact mechanism of pathogenic peptide aggregation remains a controversial topic (5).

It has been suggested that polyglutamines perturb neuronal membranes, result in their disruption concomitant with calcium dysregulation (6, 7) thereby causing Huntington’s or other amyloidogenic diseases (8-10). Huntingtin or fragments of the protein have been shown to be involved in intracellular vesicle trafficking (10-12). They associate with the ER, the Golgi-apparatus and endosomal vesicles (6, 11, 13-15). Furthermore, the length of the polyglutamine tract affects the redistribution of these polypeptides between the cytoplasm and the nucleus (16-18). Finally, mitochondrial malfunction has also been associated with the pathogenesis of Huntington’s disease (19, 20). Indeed, when fusion proteins of GST with exon 1 of huntingtin encompassing either 20 or 51 glutamines have been studied in association with brain lipid membranes protein oligomerization in the presence of long polyglutamine tracts is observed (21).

Studies on the various factors influencing the rate of aggregation include polyQ length, flanking sequences, posttranslational modification, protease activities on huntingtin and the presence of chaperones (1). On the one hand, the macromolecular assembly of huntingtin or its fragments through polyQ interactions is modulated by association of the protein with membranes (10, 22, 23). On the other hand, the length of the polyQ has an influence on these lipid interactions and the resulting membrane disruption (10, 22). Furthermore, the polyproline segment downstream of the polyQ domain has the opposite effect by reducing both the kinetics of aggregation and the formation of β-sheets by the polyQ region (22, 24).

Importantly, the first 17 amino acids of huntingtin exon 1 (htt17), directly preceding the polyQ tract, and posttranslational modifications within this region have been shown to have a strong effect on the cellular localization of huntingtin and the propensity of the protein to aggregate (6, 24-29). Thus, disease pathogenesis in transgenic mice can be inhibited by mutations of serine 13 and 16 within this htt17 domain (30). Furthermore, the htt17 sequence carries phosphorylation, SUMOlation and nuclear export sequences (1, 27, 31-35). Membrane interactions of huntingtin *in vivo* require this htt17 domain (6, 13). Furthermore, it has been shown that htt17 enhances polyglutamine oligomerization (6, 13, 23, 36-38).

Recent structural investigations reveal a highly dynamic behaviour of htt17 and the subsequent polyQ domain where htt17 association (23), its interactions with the membrane (39-43) or with other polypeptide domains are associated with random coil-helix structural transitions (44, 45). Interestingly, the htt17 and the polyQ domains mutually influence each other and their conformational properties are coupled (46, 47). More recent investigations using electron microscopy and molecular dynamics calculations reveal tadpole like structures of exon 1 where htt17 together with polyQ forms a globular head domain increasing in size as the number of glutamines increases and where the polyproline extends as the tail (48, 49). The htt17 domain plays an important role in the polyQ aggregation and *de novo*, seeded or membrane-driven mechanisms have been distinguished (23). Depending on the conditions different aggregate sizes of globular or fibrous morphologies have been described and correlated to toxicity (15, 50, 51).

Aggregation, oligomer and/or fibril formation which are the causative events for the development and progression of Huntington’s disease (15, 50, 51) require that polyglutamines are brought in contact with each other. This can occur by protein-mediated interactions of polyglutamines (1, 52, 53), by seeding or by local accumulation of polyglutamines at bilayer surfaces (23, 39, 54). Notably, the reversible membrane interactions of the amphipathic helical htt17 domain have been characterized in a quantitative and lipid-dependent manner (39) and the importance of this domain to enhance polyglutamine aggregation has been shown *in vitro* and *in vivo* (6, 13, 36-38). Another huntingtin domain mediating membrane interactions has been identified within residues 171-371, a region bearing an overall high cationic character (10). Furthermore, biochemical and cell biological assays demonstrate the potential role of these anchoring domains in disease development (10, 21, 41, 42, 55, 56).

Membrane surface induced conformational changes in proteins play a critical role in the aggregation process, for example, by concentrating and aligning polyglutamines in such a manner to promote nucleation of amyloid formation (22, 23, 39, 57, 58). Furthermore, membranes could alter aggregate morphology to specific toxic species or stabilize potentially toxic, transient aggregation intermediates (8, 59-61). Therefore, investigations of how amyloid fibrils as well as their intermediate and protofibrillar states interact with membranes is of considerable importance (22, 62). In the case of huntingtin and synuclein it has been shown that fibrils are toxic to the cells (63) and that the docking of extra-cellular aggregates to the cell membranes is a key step of the vicious propagation-amplification cycle (64). However, hardly any of these publications provide quantitative structural and biophysical data about interactions of huntingtin domains with membranes.

This prompted us to investigate in quantitative detail the role of the membrane in the polyQ association kinetics. To this end we prepared constructs involving the membrane-anchoring htt17 domain followed by polyglutamines of different length. The time-dependent structural changes were characterized in a quantitative manner using CD spectroscopy and the supramolecular complexes formed by dynamic light scattering and ThT fluorescence. By investigating htt17 in the presence of polyQ domains as short as 9 glutamines the slower aggregation kinetics allow for a more controlled, quantitative evaluation of the processes involved in aggregation and fibril formation similar to the use of htt17-polyQ constructs used in previous biophysical studies (e.g. (41, 42, 48, 56, 65)). The polyglutamine was then successively extended in small steps to quantitatively measure the effect of polyQs on the aggregation dynamics. The results reveal a pronounced dependence of the htt17-driven aggregation rates on the presence of lipid bilayers, the length of the polyglutamine tract, and polypeptide concentration.

## MATERIALS AND METHODS

### Materials

All lipids were purchased from Avanti Polar Lipids (Alabaster, AL, USA). Water (HPLC grade), acetonitrile (99.8% HPLC grade), hexafluoro-2-propanol (99.5%), trifluoroacetic acid (TFA) (99.5%) and thioflavin (ThT) were from Sigma (St. Quentin Fallavier, France).

### Peptide synthesis and purification

The peptides with the sequences shown in Table 1 were prepared by solid-phase synthesis using a Millipore 9050 automated peptide synthesizer and its standard Fmoc (9-fluorenylme-thyloxycarbonyl) chemistry. They were purified by semi-preparative reverse phase high-performance liquid chromatography (Gilson, Villiers-le-Bel, France) using a preparative C18 column (Luna, C18-100Å-5µm, Phenomenex, Le Pecq, France) and an acetonitrile/water gradient. Their identity and purity (> 90%) were verified by analytical HPLC and MALDI mass spectrometry (MALDI-TOF Autoflex, Bruker Daltonics, Bremen, Germany). The fractions of interest were lyophilized for storage at −20°C.

**Table 1:**
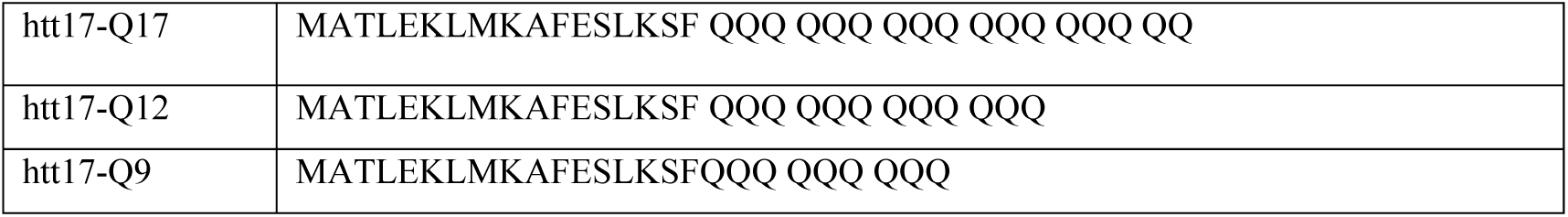
Amino acid sequences of huntingtin exon 1-related peptides.

|  |  |
| --- | --- |
| htt17-Q17 | MATLEKLMKAFESLKSF QQQ QQQ QQQ QQQ QQQ QQ |
| htt17-Q12 | MATLEKLMKAFESLKSF QQQ QQQ QQQ QQQ |
| htt17-Q9 | MATLEKLMKAFESLKSFQQQ QQQ QQQ |

### Small unilamellar vesicles

The lipids were dissolved in chloroform/methanol (2/1 v/v). The solvent was evaporated under a flow of nitrogen gas in such a manner to form a homogeneous film on the walls of glass test tubes. The remaining solvent was removed by exposure for about 12 hours to high vacuum (P < 100 mPa). Subsequently, the lipids were hydrated in 10 mM Tris-HCl, pH 7 for more than an hour and subjected to several freeze-thaw cycles. Small unilamellar vesicles (SUV) were obtained after less than 1 min of tip sonication (Bandelin Sonopuls HD 200, Berlin, Germany).

### Circular dichroism spectroscopy

Circular dichroism (CD) spectra were recorded from 260 to 194 nm (spectral resolution: 1 nm, data pitch: 1 nm, scan speed: 100 nm/min) using a J-810 spectropolarimeter (Jasco, Tokyo, Japan). 20 µL of peptide solution (at 1 mg/mL, in 10 mM Tris-HCl buffer pH 7) were transferred into a quartz cuvette of 1 mm path length and mixed with 200 µL of SUVs (C = 0.5 mg lipid/mL) to reach the final concentration of peptide ≈ 9.1·10^−2^ mg/mL (corresponding to 29, 26 and 22 μM for htt17-Q9, htt17-Q12 and htt17-Q17, respectively). The mixture was vortexed for ≈ 15 s prior to spectral acquisition. The secondary structure composition of the peptides was calculated from the spectra using the DicroProt analysis software (66).

### Measurements of thioflavin-T fluorescence

A 100 mM stock solution of Thioflavin-T (ThT) was prepared in 10 mM Tris-HCl, pH 7 and stored in the dark at −20°C. The stock solution was diluted to 1 mM in the same buffer on the day of analysis. To prepare samples for spectrofluorometry, 0.3 mg of peptide were dissolved to 0.5mg/mL in a few mL of SUV suspension and supplemented with the same volume of 10 mM Tris-HCl to reach the desired peptide-to-lipid ratio (cf. Figure captions and text). A ThT stock solution was added to the suspension to reach a final concentration of 5 µM and immediately vortexed for about one minute. 1 ml of suspension of SUVs with the peptide and the chromophore was transferred into a quartz cuvette and the sample holder of a Fluorolog 3-22 spectrometer (Horiba Jobin-Yvon, Longjumeau, France). The sample was excited at *λ*_*exc*_ = 440 nm while the dispersed fluorescence intensity was either recorded from *λ*_*fluo*_ = 455 to 651 nm or at the fixed wavelength of *λ*_*fluo*_ = 485 nm at a constant temperature of 25 °C. A resolution of Δλ = 4 nm was chosen for excitation as well as for analysis in order to ensure a good signal-to-noise ratio.

### Dynamic light scattering

Measurements were performed on a Zetasizer Nano-S system (Malvern Instruments, Worcestershire, UK) equipped with a 4 mW He-Ne laser. Samples containing the mixture of peptides and vesicles were placed in a low volume quartz cell equilibrated at 25°C, and the light scattered backward was collected at an angle of *θ* = 173°. Data analysis was performed with the DTS Malvern software implemented on a personal computer.

### Data analysis

To analyse the kinetic data, *i*.*e*. fluorescence intensity of ThT at 485 nm or circular dichroism at 208 nm as a function of time, a standard least-square fit analysis was implemented numerically. A three parameter single exponential function, *I(t) = A + B · (1 – exp (- t /* □ )) was found to describe well the intensity increase of the signals. The time constant □ was expressed in minutes.

## RESULTS

The aggregation of huntingtin exon 1-derived peptides was studied by a combination of biophysical assays. The peptides encompass the first 17-residue amphipathic sequence known to reversibly interact with membranes (39) and polyglutamine stretches of variable length (Table 1).

### CD spectroscopy

To test if membranes can increase the speed of polyglutamine aggregation and at the same time to get insight into their secondary structure, we recorded CD spectra of the polyQ peptides htt17-Q9, htt17-Q12 and htt17-Q17 as a function of time. Their structural changes where monitored in solution in the absence (Figure 1A-C) and in the presence of phospholipid vesicles (Figure 1D-F) in 10 mM Tris-HCl, pH 7 at a concentration of 0.1 mg/mL (about 30 μM). The spectra were recorded between 260 and 194 nm where the spectral line shape correlates with the secondary structure composition of the peptides. Measurements were performed every 24.5 minutes and more than 18 spectra were recorded for each polypeptide sequence.

**Figure 1:**
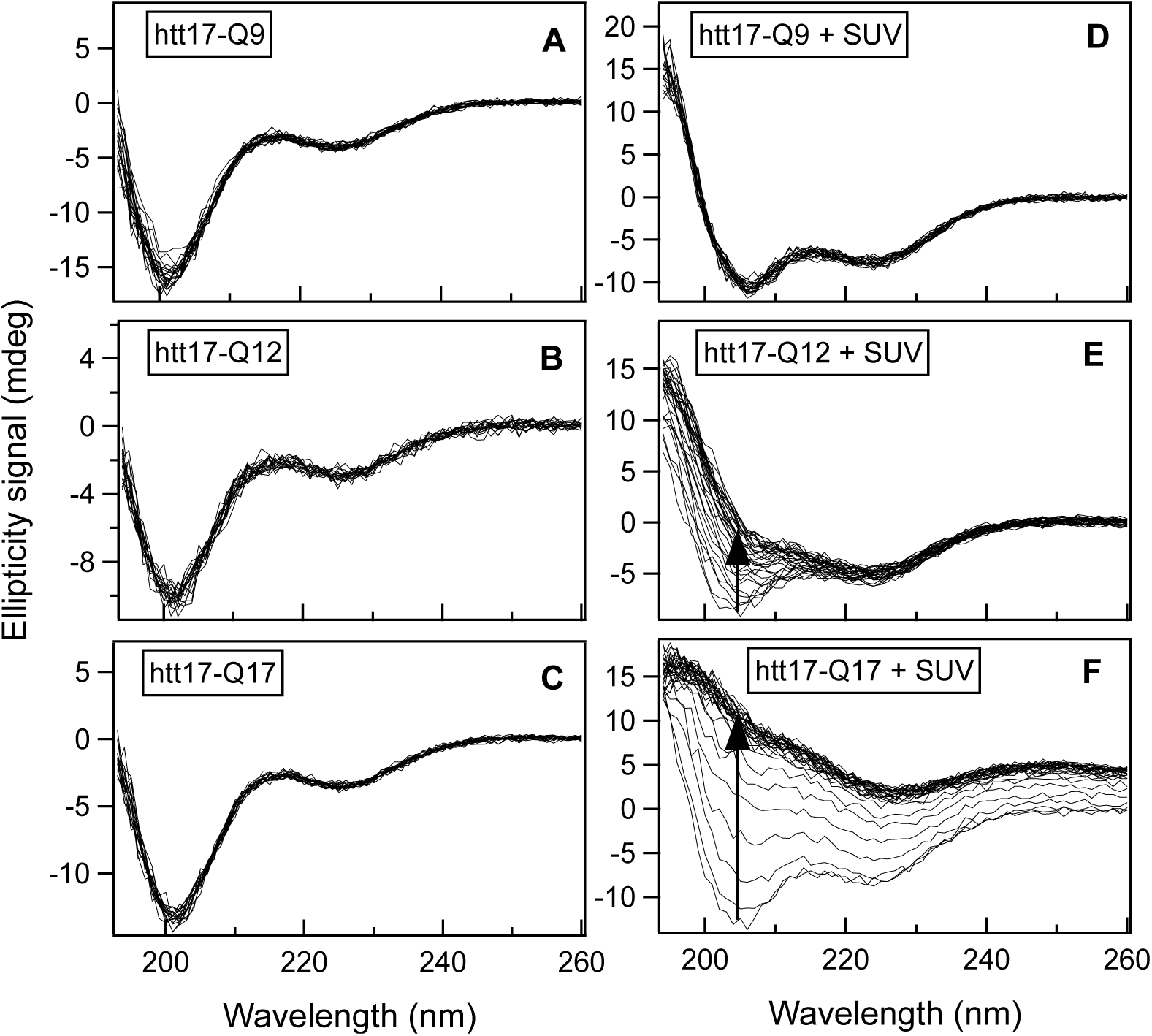
Time-dependent structural changes measured by circular dichroism. CD spectra of htt17-Q9, htt17-Q12 and htt17-Q17 (C = 9.1·10^−2^ mg/mL) in 10 mM Tris-HCl, pH 7 (A to C), and in presence of SUVs made of POPC/POPS 3/1 mole/mole (C = 0.45 mg/mL) (D to F) were recorded every 24.5 minutes. The progress of the spectral changes with time is depicted by arrows in panels E and F.

In aqueous solution the three peptides all adopt predominantly random coil conformations, without any significant spectral changes over the time period of the experiment (Figure 1A-C). When fitting the data using the DicroProt analysis software (66) less than 15% of the signal were associated with α-helical or β-sheet conformations.

Figure 1D-F exhibits the time evolution of the peptide in the presence of SUVs made of POPC/POPS at a 3/1 molar ratio. The same mass of peptides as above was mixed with a suspension of vesicles at a lipid concentration of 0.5 mg/mL in 10 mM Tris-HCl, pH 7, giving final peptide-to-lipid (P/L) molar ratios of 1/19.6, 1/22 and 1/26 for htt17-Q9, htt17-Q12 and htt17-Q17, respectively. Interestingly, the structure of htt17-Q9 remains unchanged over time since all the spectra overlap almost perfectly (Figure 1D). An estimate of the secondary structure of the peptide at the beginning of the experiment gives 25 % α-helix, 30% β-sheet and 45% random-coil structures. In contrast, the CD spectra shown in Figure 1E exhibit a significant increase in ellipticity over time in particularly between 194 and 220 nm. Whereas the initial structure of the htt17-Q12 peptide resembles that of htt17-Q9 the β-sheet content of the Q12 sequence increases gradually, at the expense of α-helix and random-coil contributions. This effect is even more pronounced for htt17-Q17 where the whole spectrum changes with time to ultimately converge to positive ellipticity values over the whole spectral range (Figure 1F).

In order to characterize the kinetics of the peptide structural changes, we quantitatively analysed the ellipticity at 208 nm over time (Figure 2). The intensity at this wavelength is related to the helix secondary structures (67). Its time dependence provides a good indication of changes in secondary structure and moreover is related to vesicle aggregation processes (68). A mono-exponential function of the form *A + B · (1 – exp (- t /* □ )), where A and B are fitting parameters and □ is the exponential time constant, describes well the intensity increase over time. The resulting fits are displayed in Figure 2 and the corresponding values of □ expressed in minutes are reported in Table 2.

**Table 2:** Time constants of the exponential increase of the circular dichroism signal measured at 208 nm (see data in Figure 2). The total peptide and lipid concentrations were kept at 9.1·10^−2^ mg/mL and 0.45 mg/mL, respectively. Thereby the peptide-to-lipid ratios are 1/19.6, 1/22 and 1/26 for htt17-Q9, htt17-Q12 and htt17-Q17, respectively.

| | P/L | $\tau$ (min) |
| --- | --- | --- |
| htt17-Q9 | 1/19.6 | 1390 |
| htt17-Q12 | 1/22 | 279 |
| htt17-Q17 | 1/26 | 28.9 |

**Figure 2:**
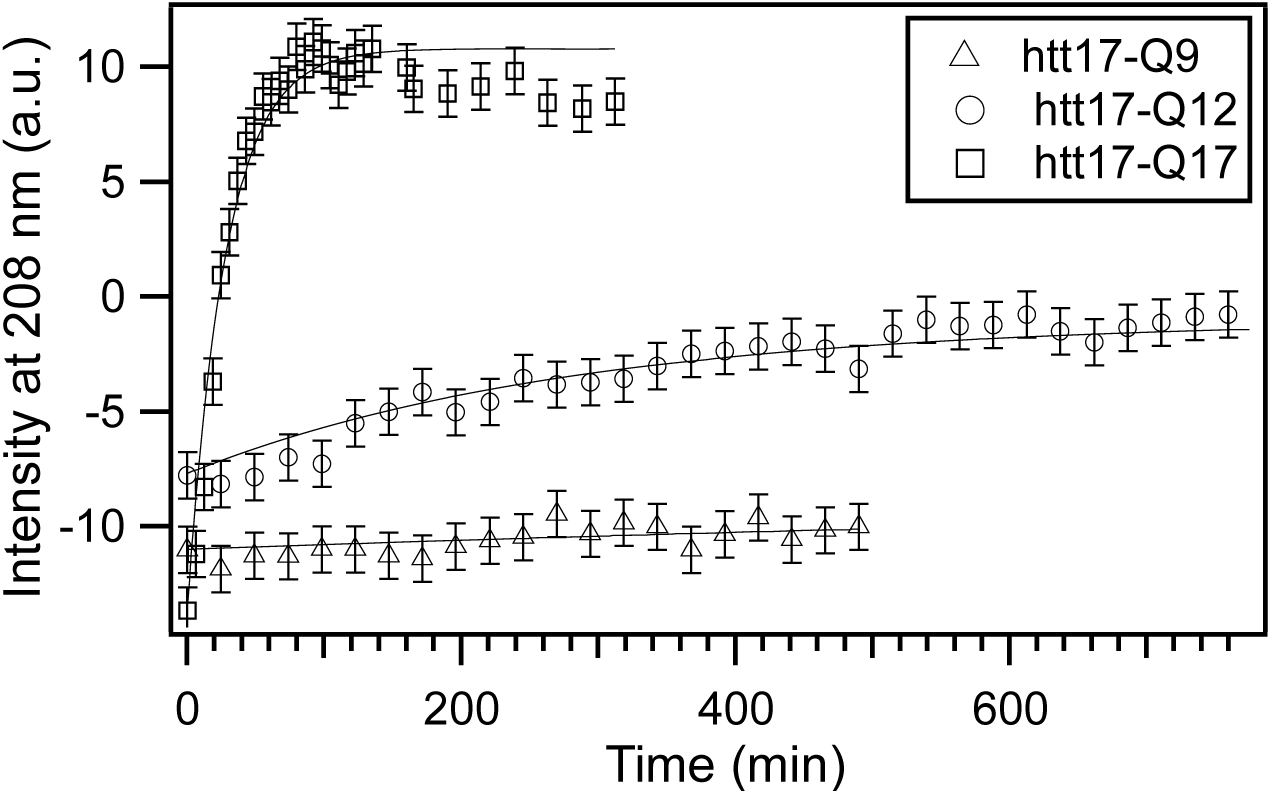
Aggregation kinetics by circular dichroism. The time-dependent intensity of the CD signal measured at 208 nm is shown for htt17-Q9, htt17-Q12 and htt17-Q17 in the presence of 0.45 mg/mL SUVs made of POPC/POPS 3/1 mole/mole in 10 mM Tris-HCl, pH 7. The results of least squares fits with a mono-exponential function are displayed as solid lines. The corresponding values of the time constants are reported in Table 2.

### Thioflavin T fluorescence

To follow the kinetics of htt17-polyQ β-sheet formation changes in the Thioflavin T (ThT) fluorescence were monitored in the presence of phospholipid bilayers. ThT is a popular reporter of amyloid aggregation because it demonstrates a strong shift and enhanced intensity of fluorescence emission upon binding to β-sheet rich fibrils (69, 70). The dye has been used to visualize and quantify the presence of misfolded protein or peptide aggregates *in vitro* and *in vivo* (71).

Figure 3 displays three sets of measurements performed at the same ThT concentration. The intensity of the fluorescence emission spectra in 10 mM Tris-HCl buffer at pH 7 increases with time in the presence of both ≈14.5 µM htt17-Q17 and SUVs made of POPC/POPS 3/1 (Figure 3A). In contrast, the fluorescence measured in control experiments with peptides only (Fig. 3B) or with SUVs only (Fig. 3C) remains unchanged.

**Figure 3:**
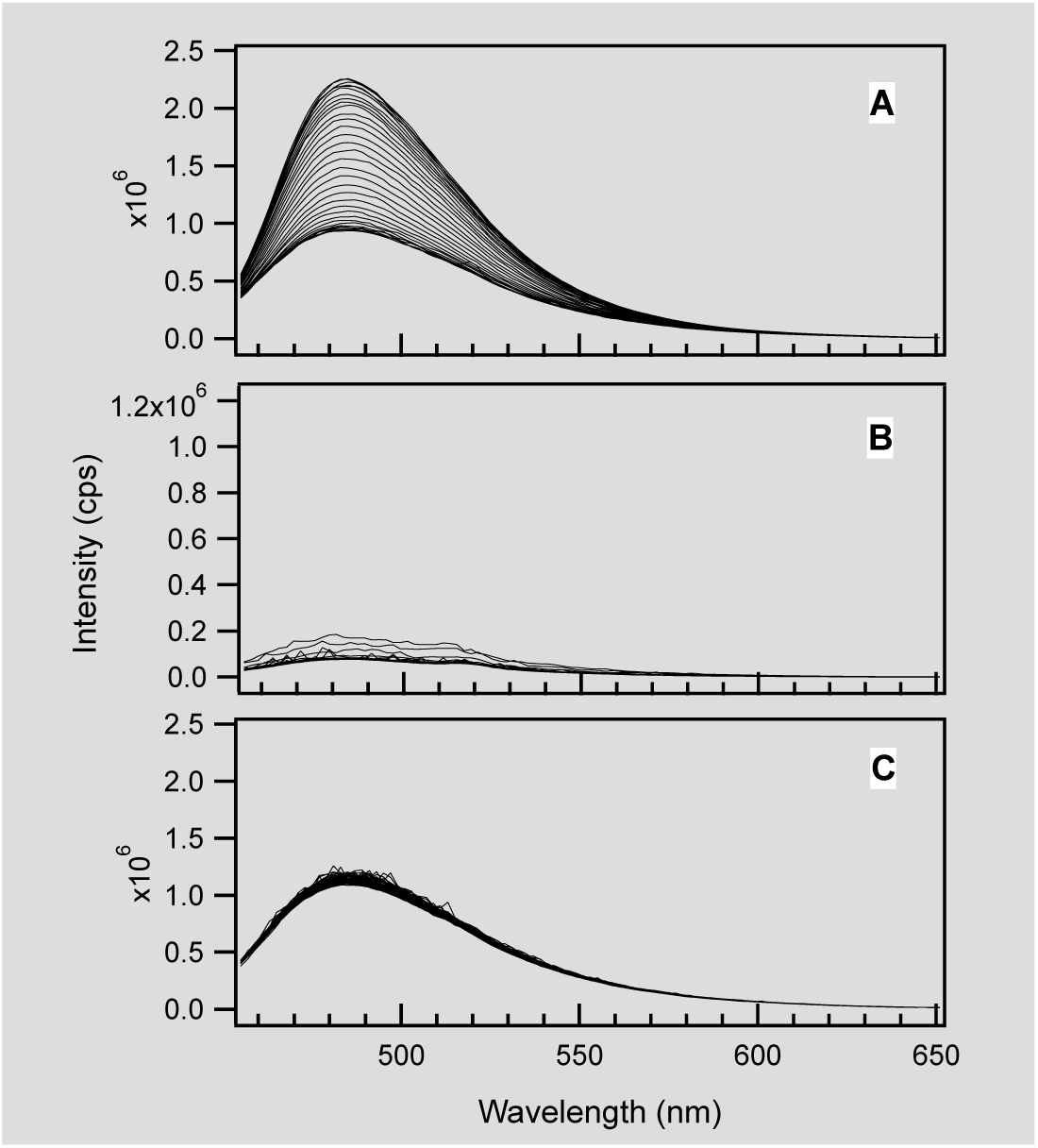
Time-dependent amyloid formation of htt17-Q17. The thioflavin T fluorescence increase over time is shown in the presence of 14.5 µM htt17-Q17 and SUVs made of 320 µM POPC/POPS 3/1 mole/mole in 10 mM Tris-HCl buffer, pH 7 (A). The peptide-to-lipid molar ratio was 1/22. The control experiments show almost no changes in thioflavin T florescence when exposed to the same amount of htt17-Q17 only (B) or SUVs only (C). The thioflavin T concentration was 5 µM in all recordings.

In order to characterize the kinetics of β-sheet formation in a quantitative manner, we measured the fluorescence intensity of ThT in the presence of the three peptides over time at the fixed wavelength of *λ*_*fluo*_ = 485 nm, while the dye was continuously excited at *λ*_*exc*._ = 440 nm. To quantify the effect of increasing peptide concentrations, for each htt17-polyQ sequence three measurements were performed, namely at P/L ratios of 1/88, 1/44 and 1/22. Under most experimental conditions, an increase of the signal was measured over time, indicating peptide aggregation. In some cases, the signal first increased and then decayed, which is probably due to sedimentation of the peptides, aggregated and/or associated with the vesicles (72).

As illustrated in Figure 4, the P/L ratio has a direct effect on the efficiency of the aggregation process as well as on its kinetics. For all three peptides, increasing the P/L ratio makes the aggregation process more efficient. Therefore, the fluorescence intensities measured at L/P=22 are higher than for P/L=1/44 or P/L=1/88.

**Figure 4:**
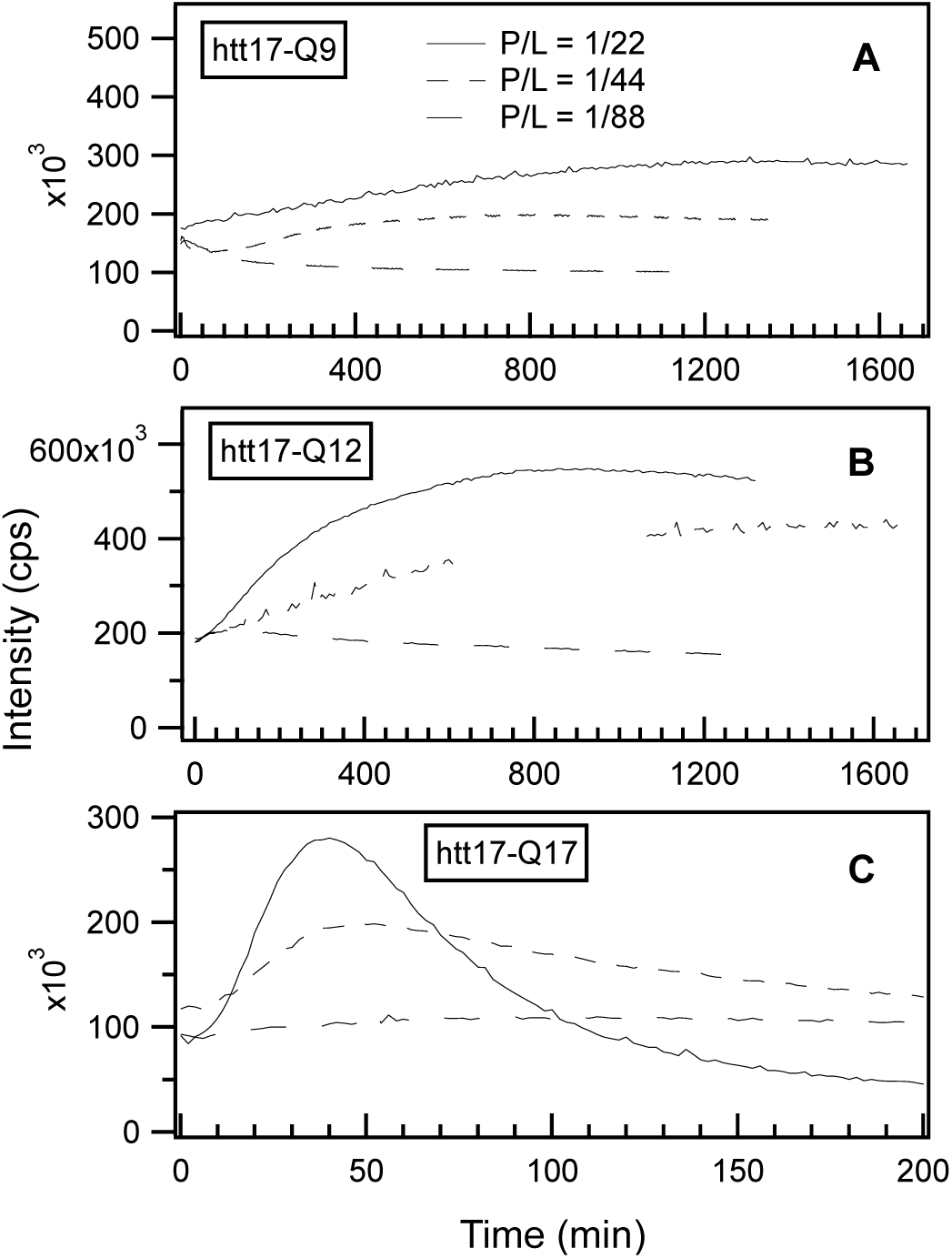
Time-dependent amyloid formation as a function of polypeptide concentration. The thioflavin T fluorescence was measured at 485 nm in the presence of SUVs and htt17-Q9 (A), htt17-Q12 (B) or htt17-Q17 (C), in 10 mM Tris-HCl, pH7. The peptide-to-lipid molar ratios are 1/22, 1/44 and 1/88, displayed as solid, short-dashed and long-dashed lines, respectively. The lipid concentration was kept constant (C ≈ 320 µM) while the amount of peptide was adjusted to obtain the P/L ratios indicated. The ThT concentration was ≈ 5 µM in all the recordings.

To quantitatively compare the kinetics of β-sheet formation, we used the same model as for the treatment of the CD data (cf. above). The mono-exponential function *A + B · (1 – exp (- t /* □ )) was used to fit the signal increase and the resulting exponential time constants □ are reported in Table 3. For htt17-Q9 at the P/L ratios 1/44 and 1/88, and for htt17-Q12 at P/L=1/88 the signals exhibit slightly negative slopes at the beginning of the experiments. Therefore, these data were not analyzed in a quantitative manner.

**Table 3:**
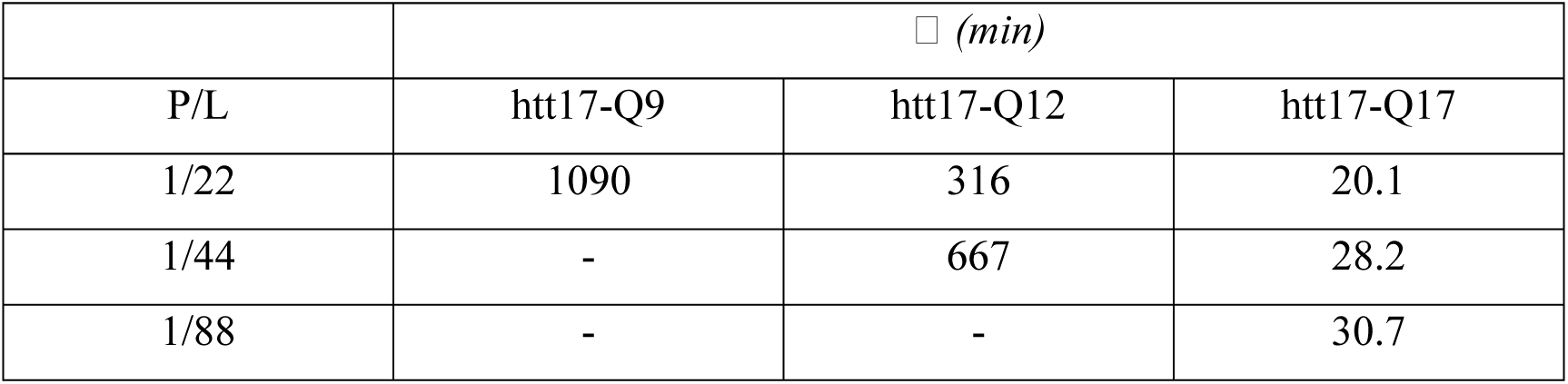
Time constants of the exponential increase of fluorescence intensity of thioflavin T in the presence of SUVs and htt17-Q9, ht17-Q12 or htt17-Q17 (*see* Figure 4). The lipid concentration was kept constant (C ≈ 320 µM) and the peptide concentrations were adjusted to obtain the desired peptide-to-lipid ratios of 1/22, 1/44 or 1/88.

| | $\tau$ (min) | | |
| --- | --- | --- | --- |
| P/L | htt17-Q9 | htt17-Q12 | htt17-Q17 |
| 1/22 | 1090 | 316 | 20.1 |
| 1/44 | - | 667 | 28.2 |
| 1/88 | - | - | 30.7 |

### Dynamic light scattering

To get a more detailed view of the vesicle-vesicle interactions and the subsequent sedimentation processes that was suspected to occur in some of the fluorescence and CD spectroscopy experiments, we performed dynamic light scattering measurements (DLS) where the three htt17-polyQ peptides were exposed to suspensions of SUVs made of POPC/POPS 3/1 mole/mole (0.45 mg/mL, i.e. 580 μM) in 10 mM Tris-HCl, pH 7 buffer (Figure 5). In a first series of experiments the htt17-Q9 concentration was 29 μM. The hydrodynamic diameter of the systems and their corresponding polydispersity indexes (PDI), were measured every 18.7 minutes for more than 7 hours. Related experiments were performed with 25.9 μM htt17-Q12 and 22.9 μM htt17-Q17, respectively, i.e. the same concentrations by weight. This corresponds to the same experimental conditions also used for the CD measurements. When vesicles and htt17-Q17 are mixed the DLS data indicate a strong and continuous increase in hydrodynamic radius over the whole time period, while the htt17-Q9 peptide left the light scattering unaffected. Thereby, the apparent changes in ellipticity were assigned to light scattering processes induced by supra-wavelength sized systems like vesicle and/or peptide clusters.

**Figure 5:**
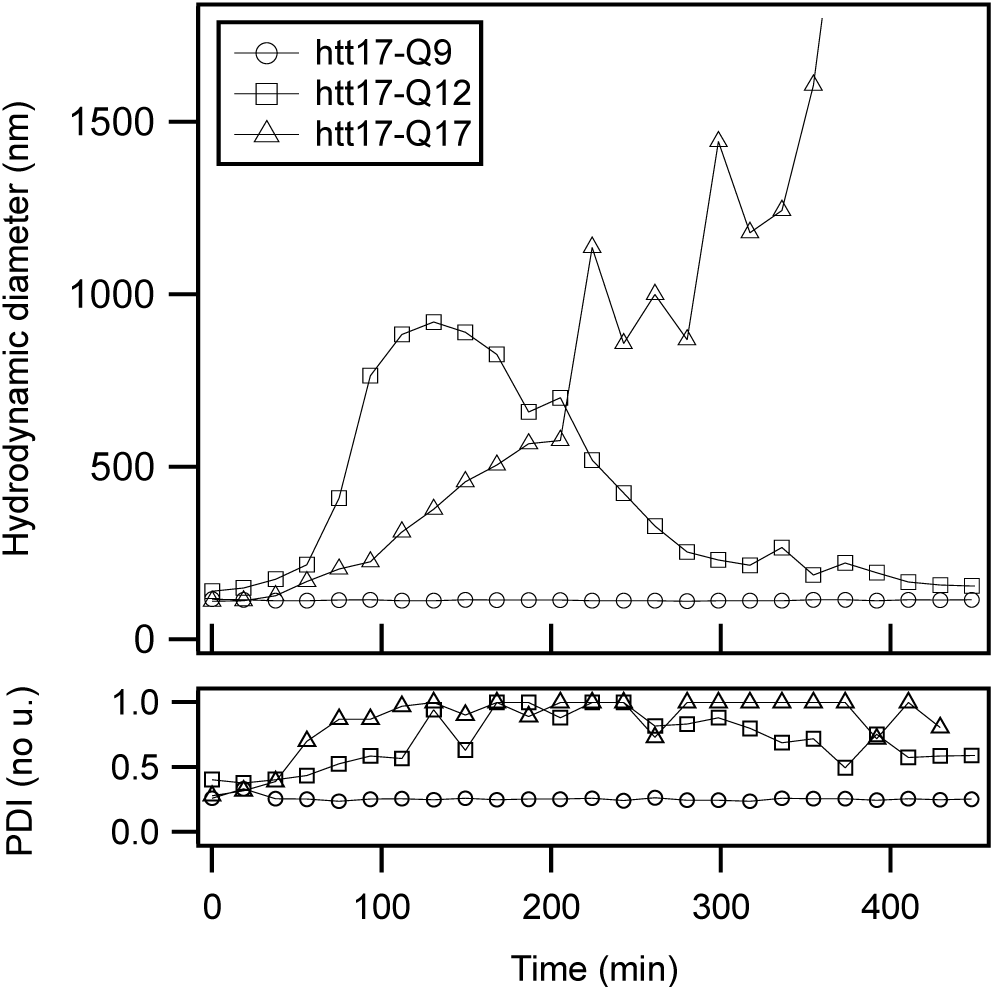
Time-dependent changes of aggregate size. Hydrodynamic diameter as a function of time of SUVs in the presence of htt17-Q9 (circles), htt17-Q12 (squares) or htt17-Q17 (triangles) in 10 mM Tris-HCl, pH7. Measurements were performed by dynamic light scattering in the absence of mechanical stirring. The bottom frame shows the polydispersity index (PDI) of the corresponding measurements. The same concentrations of lipids and peptides were used as in the CD experiments presented in Figures 1 and 2.

## DISCUSSION

Previously it has been shown that huntingtin carries domains before and after the poly-Q tract that promote its reversible membrane association and it has been suggested that this membrane-association helps in polyQ aggregation (10, 23, 26, 39). In particular, the N-terminal 17 amino acids preceding the polyQ tract of huntingtin (usually abbreviated htt17 or N17) are involved in the regulation of the spatio-temporal distribution of huntingtin or fragments thereof. Their association with membrane components and with different cellular compartments has been shown important for the development of symptoms of Huntington’s disease (6, 10, 13, 16–20, 26, 30, 36–38). Here we show that htt17-polyQ membrane association indeed strongly accelerates polypeptide aggregation in a manner that is dependent on the number of glutamines. A series of sequences was investigated with htt17 as a membrane anchor and various polyQ extensions (Table 1).

In order to quantitatively measure the aggregation kinetics biophysical experiments were performed at a polypeptide concentration of about 30 μM. This relatively low concentration is still suitable for spectroscopic analysis, but longer polyQ constructs tend to aggregate at very short time scales. Therefore, quantitative studies of the speed of aggregation were performed with constructs carrying a limiting number of glutamines (*see* also (73)). It should be noted that in their natural context the solubility of the full-length protein or its exon 1 domain is increased by the polyproline flanking sequence following the polyQ domain (28, 29, 37, 44). Furthermore, in the cellular environment, interactions with other proteins, chaperones and proteases, as well as other domains of the full-length protein assure that healthy cells are protected from huntingtin aggregation (74, 75). Therefore, it takes many years to develop the disease even when much longer polyQ mutants are present (1).

When diluted in aqueous buffer CD spectra of the peptides with short polyQ additions are indicative of predominantly random coil structure and some features of α-helical/β-sheet conformations. Upon addition of POPC/POPS 3/1 membranes the CD spectra of htt17-polyglutamines carrying 12 or more glutamines change in appearance over the next few hours (Fig 1). The increase in hydrodynamic radius observed by DLS (Fig. 5) and by thioflavin fluorescence (Figs. 3 and 4) are indicative that the peptides aggregate in the presence of membranes in a manner that depends on the peptide-to-lipid ratio (Fig. 4). In the case of htt17-Q17 the changes in the CD spectra obtained at concentrations of about 30 μM peptide and 650 μM lipid reach saturation after about 1.5h (Figs. 1 and 2).

At longer incubation times CD- and ThT fluorescence spectra decrease in intensity (Figs. 2 and 4C) probably because the larger peptide-lipid aggregates sink to the ground and/or stick to the surface of the glass tubes. The spectral changes are suggestive of an increase in β-sheet conformation (Fig. 1) in agreement with other structural data (45, 65), when at the same time a continuous increase in hydrodynamic radius is observed (Fig 5). Concomitant with this aggregation a quantitative analysis of the resulting CD spectra is hampered by light diffraction artefacts (Figs. 1F, 2 and 5). Peptides carrying a shorter polyQ segment aggregate more slowly or do not show spectral changes at all (Fig 1D,E, Table 2). The data summarized in Table 2 indicates that not only the degree of aggregation but also its kinetics accelerate with an increasing number of glutamines.

Whereas under the conditions investigated the peptides studied in this work remain in solution over many hours, even days in the absence of POPC/POPS vesicles, here we show that the addition of membranes strongly catalyses the aggregation process. Indeed, whereas aggregate formation *de novo* or through seeding are well established pathways, membranes have been shown to provide a third aggregation mechanism (23). The data presented is in agreement with previous investigations where GDT-exon1 constructs have been found associated with rat postsynaptic membranes and where brain lipid vesicles accelerate nucleation and thereby fibril formation upon trypsin cleavage of 3 μM GST-exon 1 encompassing 51 glutamines (21). In the same study the presence of zwitterionic DMPC or of DOPC/SM/cholesterol slowed down this effect thereby being in-line with studies where htt17 membrane interactions were weak or absent for zwitterionic and cholesterol-rich membranes (39, 76).

The N-terminal 17-residue sequence has been demonstrated to adopt a largely helical conformation when being membranes-associated (41, 42, 77, 78), when part of a htt17-polyQ fiber (45, 46, 79) or in aggregation intermediates (51, 73). Here we have shown that the membrane interactions of the htt17 flanking region of the polyQ domain promotes the aggregation process of huntingtin exon 1. Htt17 also plays a leading role in catalysing the nucleation process during polyQ aggregation in solution. The htt17 amphipathic helix has been shown to associate into small oligomeric structures which serve as nucleation sites (23, 51, 73) from which polyQ fibrils elongate (79, 80). Furthermore, the flanking sequences play important additional roles in regulating the proteolytic degradation of the protein and its aggregates where the toxicity depends in a complex manner on the conformational subpopulation rather than the polyQ aggregation propensity (27, 29, 51, 81).

Flanking regions of other polyQ proteins seem equally important in the regulation of their aggregation and fiber formation (1). In particular previous studies showed that posttranslational modifications of flanking regions such as phosphorylation, SUMOylation or ubiquination including of htt17 have an effect on polyQ aggregation (1, 27, 31–34). This has been attributed to the changes of htt17 oligomerization being a consequence of such modifications, as well as modifications of its interaction surface with chaperones. The work presented here suggest that also its interactions with membranes should be strongly modified when e.g. negative phosphates are attached to its serines (25), when lysines are made unavailable for protein-protein and protein-lipid interactions (77, 78, 82) or the overall positive charge of htt17 is neutralized or inverted by posttranslational modifications thus abolishing the electrostatic attraction of the flanking regions to anionic membranes (10, 77). The numerous possibilities to interfere with protein-protein and protein-lipid interactions and thereby fiber formation explain why not all polyQ-extension diseases follow the same pattern and why the age onset of Huntington disease shows significant variation even for the same number of glutamines (1).

We suggest that the amphipathic htt17 helix, which reversibly associates with the membrane interface (77, 78), concentrates and aligns the polyQ chains in such a manner to facilitate intermolecular interactions and fiber association (10, 39). Once the interactions with the membrane are established the aggregation process is enhanced by high peptide density (i.e. P/L ratio) and by extended polyQ chains (Tables 2 and 3) but slowed down by the presence of the polyproline stretch that follows the poly-Q domain (44, 83). Thereby these and other biophysical observations provide a rationale for a number of biochemical and cell biological experiments where huntingtin has been shown associated with membranes of intracellular organelles (6, 10, 13, 16–20, 25) or where the membrane anchoring domain htt17 has been demonstrated to promote the development of the disease (6, 13, 26, 30, 36–38).

Such lipid interactions have been shown to be modulators of aggregation and fibril formation also for α-synuclein (57, 62, 64), islet amyloid polypeptide (9) and β-amyloid (58, 84). Within these studies the detailed membrane composition including the resulting physico-chemical properties such as negative charge density, fluidity, saturation, curvature or interactions with specific lipids all play important roles in the aggregation process (9, 39, 56, 76, 85). Thereby, the membrane interactions of polyQ flanking regions and their modulation by posttranslational modifications provides a possible therapeutic intervention site which has to our knowledge not been explored in greater detail.

## Abbreviations used

CD: circular dichroism
DLS: dynamic light scattering
HD: Huntington’s disease
HFIP: 1,1,1,3,3,3-hexafluoro-2-propanol
HPLC: high-performance liquid chromatography
MALDI: matrix-assisted laser desorption ionization
MLV: multilamellar vesicles
SUV: small unilamellar vesicles
POPC: 1-palmitoyl-2-oleoyl-*sn*-glycero-3-phosphocholine
POPS: 1-palmitoyl-2-oleoyl-*sn*-glycero-3-phosphoserine
TFA: trifluoroacetic acid
ThT: thioflavine T
TOF: time-of-flight

## ACKNOWLEDGEMENTS

We are grateful to Nicole Harmouche, Rabia Sarroukh, Caroline Lonez, Erick Goormaghtigh and Jean-Marie Ruysschaert for providing data of unpublished experiments and for discussion as well as Delphine Hatey for technical help during peptide synthesis and purification. The financial contributions of the Agence Nationale de la Recherche (projects MemPepSyn 14-CE34-0001-01, InMembrane 15-CE11-0017-01, Biosupramol 17-CE18-0033-3 and the LabEx Chemistry of Complex Systems 10-LABX-0026_CSC), the University of Strasbourg, the CNRS, the Région Grand-Est and the RTRA International Center of Frontier Research in Chemistry, the American Foundation for Research on Huntington’s Disease (CHDI) are gratefully acknowledged.

## Author contributions

AM performed and analysed the experiments and helped in writing the paper. BB assured funding, designed experiments, discussed data and wrote the paper.

